# Functional evaluation of the P681H mutation on the proteolytic activation the SARS-CoV-2 variant B.1.1.7 (Alpha) spike

**DOI:** 10.1101/2021.04.06.438731

**Authors:** Bailey Lubinski, Maureen H. V. Fernandes, Laura Frazier, Tiffany Tang, Susan Daniel, Diego G. Diel, Javier A. Jaimes, Gary R. Whittaker

## Abstract

Severe acute respiratory syndrome coronavirus 2 (SARS-CoV-2) is the agent causing the COVID-19 pandemic. SARS-CoV-2 B.1.1.7 (Alpha), a WHO variant of concern (VOC) first identified in the UK in late 2020, contains several mutations including P681H in the spike S1/S2 cleavage site, which is predicted to increase cleavage by furin, potentially impacting the viral cell entry. Here, we studied the role of the P681H mutation in B.1.1.7 cell entry. We performed assays using fluorogenic peptides mimicking the Wuhan-Hu-1 and B.1.1.7 S1/S2 sequence and observed no significant difference in furin cleavage. Functional assays using pseudoparticles harboring SARS-CoV-2 spikes and cell-to-cell fusion assays demonstrated no differences between Wuhan-Hu-1, B.1.1.7 or a P681H point mutant. Likewise, we observed no differences in viral growth between USA-WA1/2020 and a B.1.1.7 isolate in cell culture. Our findings suggest that while the B.1.1.7 P681H mutation may slightly increase S1/S2 cleavage this does not significantly impact viral entry or cell-cell spread.

**Highlights:**

- SARS-CoV-2 B.1.1.7 VOC has a P681H mutation in the spike that is predicted to enhance viral infection
- P681H does not significantly impact furin cleavage, viral entry or cell-cell spread
- Other mutations in the SARS-CoV-2 B.1.1.7 VOC may account for increased infection rates

**Graphical abstract:** 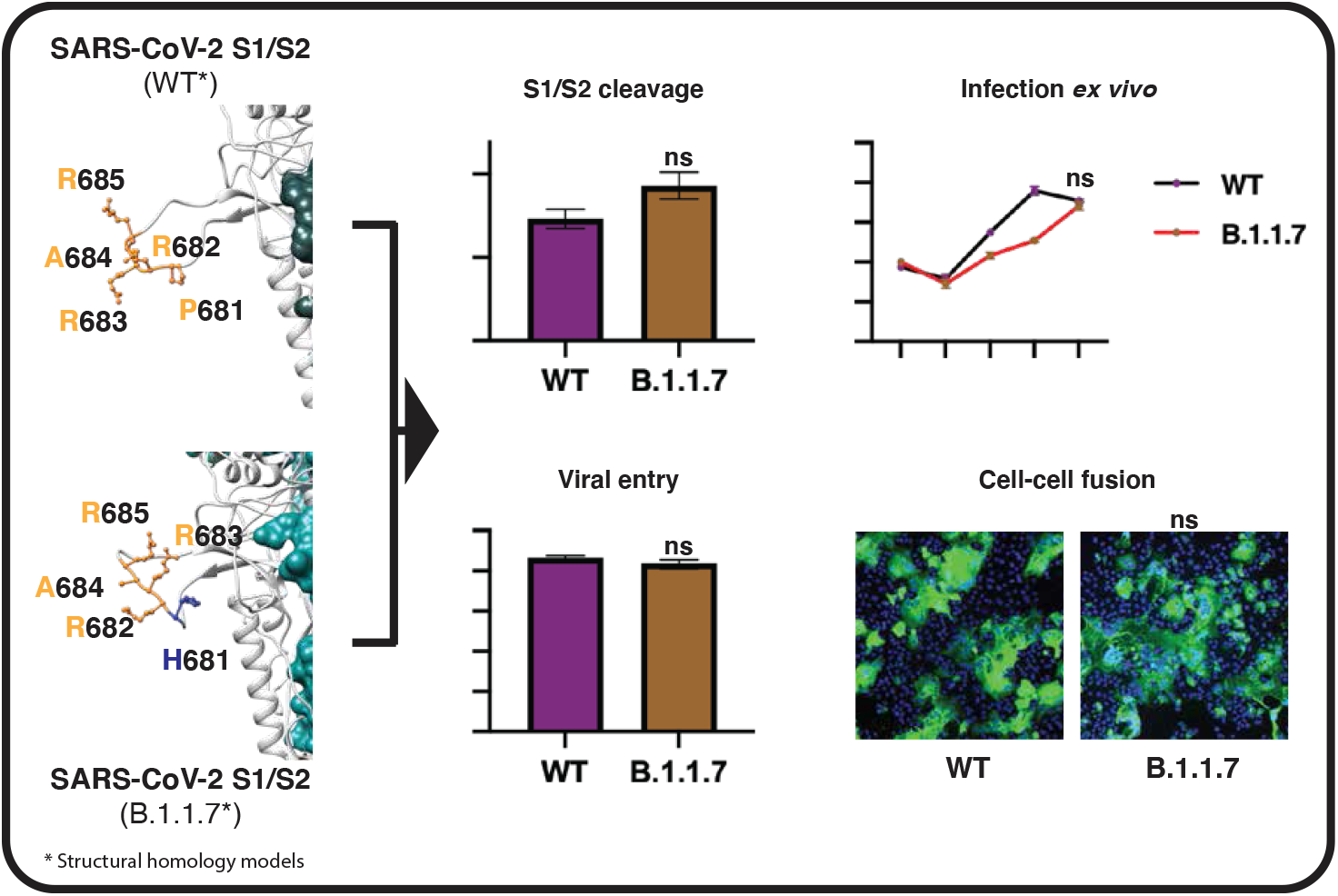

## Introduction

Severe acute respiratory syndrome coronavirus 2 (SARS-CoV-2) is the agent behind the current COVID-19 pandemic (Whittaker et al., 2021). SARS-CoV-2 emerged from a yet to be determined animal reservoir and was first identified in late 2019; it has since rapidly spread throughout the world. The virus now exists in two lineages, A and B. While both lineages remain in circulation globally, the B lineage became the dominant virus following its introduction into Northern Italy in February 2020. The B lineage has undergone significant diversification as it expanded, and in particular acquired an S gene mutation (D614G) that resulted in a more stabilized spike protein, which has been linked to increased transmissibility (Zhou et al., 2021). D614G has now become established in circulating B lineage viruses.

In late summer to early fall 2020, a new variant of SARS-CoV-2 was identified in the UK, based on S-gene target failures in community-based diagnostic PCR testing (Rambaut et al., 2020). Following subsequent sequence analysis, this variant was defined as variant of concern (VOC) B.1.1.7/501Y.V1 by Public Health England, and later denominated as the Alpha variant. B.1.1.7 rapidly expanded across the south of England and subsequently spread to other parts of the U.K. (Volz et al., 2021), and then globally. The B.1.1.7 VOC was unusual compared to other SARS-CoV-2 variants emerging at that time in that is contained a higher than typical level of point mutants across its genome; 23 in total. Of particular note were nine mutations in the spike gene compared to prototype sequences; a 69-70 deletion, Y144 del, N501Y, A570D, D614G, P681H, T716I, S982A, and D118H, with seven of these distinct to B.1.1.7. One of the more concerning mutations was N501Y, which was linked (along with D614G) to increased affinity of the spike protein to the SARS-CoV-2 receptor, angiotensin converting enzyme 2 (ACE2). N501Y has subsequently been found in other VOCs circulating around the world, i.e., B.1.351 and B.1.1.28.1 (P.1) (Coutinho et al., 2021; Lauring and Hodcroft, 2021; Tegally et al., 2021a).

Since its first identification, SARS-CoV-2 B.1.1.7 has undergone extensive characterization. Current consensus indicates that this VOC has a transmission advantage in the community (Davies et al., 2021; Volz et al., 2021), possibly accompanied by increased disease severity (Challen et al., 2021), but it does not appear to evade immune surveillance by natural immunity or vaccination. One hypothesis is that the B.1.1.7 variant acquired its extensive range of mutations in a single immunocompromised individual (Rambaut et al., 2020) who then initiated a super-spreader event that gave rise to the subsequent dissemination of the lineage.

The P681H mutation of B.1.1.7 is of note as it is part of a proteolytic cleavage site for furin and furin-like proteases at the junction of the spike protein receptor-binding (S1) and fusion (S2) domains (Jaimes et al., 2020a). The S1/S2 junction of the SARS-CoV-2 S gene has a distinct indel compared to all other SARS-like viruses (*Sarbecoviruses* in *Betacoronavirus* lineage B) the amino acid sequence of SARS-CoV-2 S protein is _681_-P-R-R-A-R|S-_686_ with proteolytic cleavage (|) predicted to occur between the arginine and serine residues depicted. Based on nomenclature established for proteolytic events (Polgár, 1989), the R|S residues are defined as the P1|P1’ residues for enzymatic cleavage, with 681H of B.1.1.7 S being the P5 cleavage position. The ubiquitously-expressed serine protease furin is highly specific and cleaves at a distinct multi-basic motif containing paired arginine (R) residues; furin requires a minimal motif of **R**-X-X-**R** (P4-X-X-P1), with a preference for an additional basic residue (histidine – H or lysine – K) at P2; i.e., **R**-X-**K/H**-**R** (Seidah and Prat, 2012). For SARS-CoV-2, the presence of the S1/S2 “furin site” enhances virus transmissibility (Johnson et al., 2021; Peacock et al., 2020). For B.1.1.7 S, P681H (P5) may provide an additional basic residue (especially at low pH) and modulate S1/S2 cleavability by furin, and hence virus infection properties.

We previously studied the role of proteolytic activation of the spike protein of the prototype lineage B SARS-CoV-2 (isolate Wuhan-Hu-1) (Tang et al., 2021). Here, we used a similar approach to study the role of the proteolytic activation of the spike protein in the context of the B.1.1.7 VOC, with a focus on the P681H point mutant.

## Results

### B**ioinformatic and biochemical analysis of the SARS-CoV-2 B.1.1.7 S1/S2 cleavage site**

To gain insight into proteolytic processing at the S1/S2 site of the B.1.1.7 spike protein, we first took a bioinformatic approach utilizing the PiTou (Tian et al., 2012) and ProP (Duckert et al., 2004) cleavage prediction tools, comparing B.1.1.7 to the prototype virus Wuhan-Hu-1 as well as to SARS-CoV, MERS-CoV, and selected other human respiratory betacoronaviruses (HCoV-HKU1 and HCoV-OC43) (Figure 1A). Both algorithms predicted a limited increase in the furin cleavage for B.1.1.7 compared to Wuhan-Hu-1; in comparison SARS-CoV is not predicted to be furin-cleaved; as expected, MERS-CoV showed a relatively low furin cleavage score with HCoV-HKU1 and HCoV-OC43 showing much higher furin cleavage scores. Previously, we showed that the SARS-CoV-2 S1/S2 cleavage site is predicted to fold as a flexible loop exposed from the spike structure (Jaimes et al., 2020a). Considering that the P681H mutation was predicted to slightly increase the furin cleavage at this site, we modeled the SARS-CoV-2 B.1.1.7 variant spike and observed no marked changes in the predicted folding, compared to Wuhan-Hu-1 (Figure 1B).

**Figure 1.**
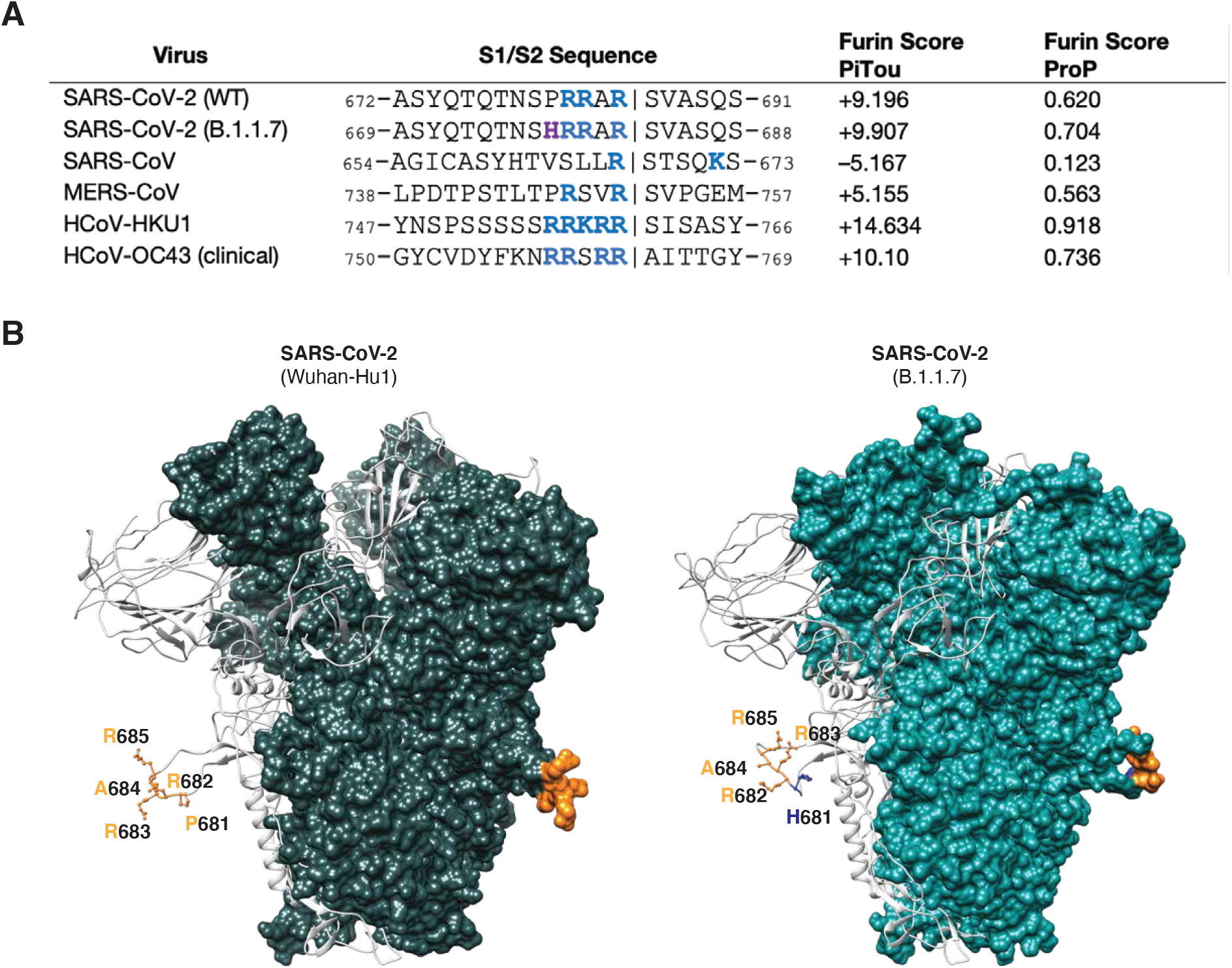
SARS-CoV-2 spike S1/S2 cleavage site. **A**. Furin cleavage score analysis of CoV S1/S2 cleavage sites. CoV S sequences were analyzed using the ProP 1.0 and PiTou 3.0 furin prediction algorithm, generating a score with bold numbers indicating predicted furin cleavage. (|) denotes the position of the predicted S1/S2 cleavage site. Basic resides, arginine (R) and lysine (K), are highlighted in blue, with histidine in purple. Sequences corresponding to the S1/S2 region of SARS-CoV-2 (QHD43416.1), SARS-CoV (AAT74874.1), MERS-CoV (AFS88936.1), HCoV-HKU1 (AAT98580.1), HCoV-OC43 (KY369907.1) were obtained from GenBank. Sequences corresponding to the S1/S2 region of SARS-CoV-2 B.1.1.7 (EPI_ISL_1374509) was obtained from GISAID. **B**. SARS-CoV-2 spike structural models. Homology models were built for the spike protein from Wuhan-Hu1 (WT) and B.1.1.7 variants. The S1/S2 cleavage site sequences are noted. WT sequence _681_PRRAR_685_ is noted in orange and mutated residue H681 is noted in blue.

To directly address the activity of furin on the SARS-CoV-2 B.1.1.7 S1/S2 site, we used a biochemical peptide cleavage assay (Jaimes et al., 2019). The specific peptide sequences used here were TNSHRRARSVA (B.1.1.7 S1/S2) and TNSPRRARSVA (Wuhan-Hu-1 S1/S2). We tested furin, along with trypsin as a control as described previously (Jaimes et al., 2020b). We also assessed the effect of lowered pH because of the known properties of histidine (H) to have an ionizable side chain with a p*K*a near neutrality (Nelson and Cox, 2000) (Figure 2A). As predicted, furin effectively cleaved Wuhan-Hu-1 (WT) S1/S2, with B.1.1.7 S1/S2 showing a slight increase in cleavage at pH 7.5. At pH 7.0 and pH 6.5, B.1.1.7 S1/S2 was actually less efficiently cleaved than WT, with cleavage not measurable below pH 6.5 (data not shown). Trypsin cleaved both peptides, but was less efficient than furin (at pH 7.4). This comparative data with SARS-CoV S1/S2 sites reveals that the acquisition of the 681H mutation does not significantly increase cleavability by furin and may in fact be inhibitory at lowered pH.

**Figure 2.**
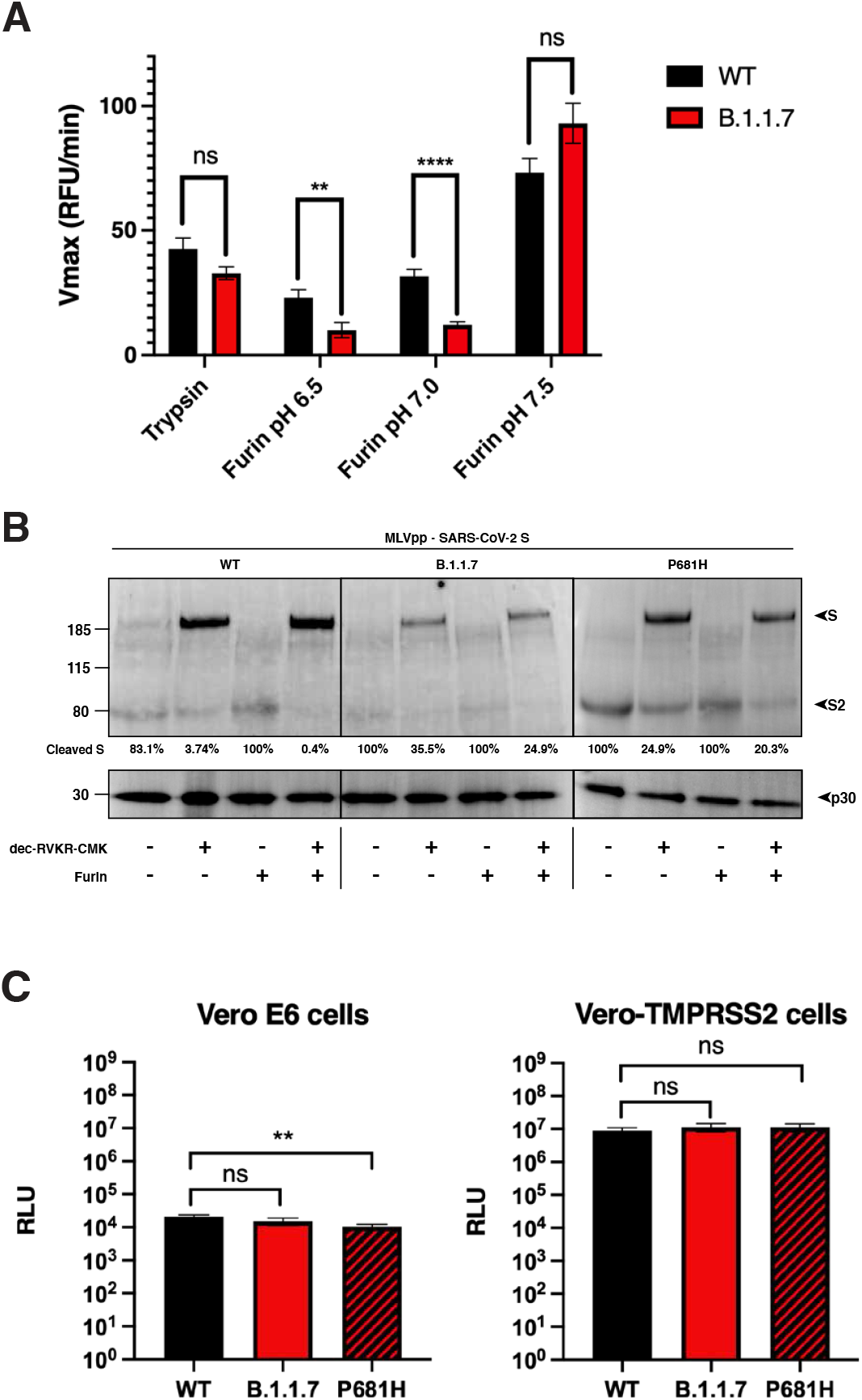
SARS-CoV-2 B.1.1.7 variant S1/S2 cleavage site activation and role in viral entry. **A**. Fluorogenic peptide cleavage assays of the SARS-CoV-2 S1/S2 cleavage site. Peptides mimicking the S1/S2 site of the SARS-CoV-2 WT and B.1.1.7 variants were evaluated for *in vitro* cleavage by trypsin and furin proteases at pH 7.4 (trypsin), and pH 6.5, 6.0 and 7.5 (furin) conditions. A significant decrease in the cleavage of the B.1.1.7 S1/S2 peptide by furin was observed at pH 6.5 and 7.0 compared to WT. In contrast, a non-significant increase in the furin cleavage of the B.1.1.7 peptide was observed at pH 7.5. **B**. Western blot analysis of MLV pseudoparticles (MLVpps) carrying the WT, B.1.1.7, or P681H S. Uncleaved S (∼185 kDa) and cleaved S2 (∼85 kDa) detection was conducted using an antibody targeting the SARS-CoV-2 S2 domain. + dec-RVKR-CMK refers to MLVpp produced in HEK-293T cells treated with 75 μM dec-RVKR-CMK at the time of transfection. + furin refers to particles treated with 6 U of recombinant furin for 3 h at 37 °C. **C**. Ratio of the intensity of cleaved S band to uncleaved S in MLV pseudoparticles. Band intensity was normalized to the uncleaved band intensity of each pseudoparticle and cleavage ratio was calculated. **D**. Pseudoparticle infectivity assays in Vero E6 and Vero-TMPRSS2 cells. Cells were infected with MLV pseudoparticles harboring the VSV-G, SARS-CoV-2 S WT, SARS-CoV-2 S B.1.1.7 variant, SARS-CoV-2 S WT with P681H mutation. Data represents the average luciferase activity of cells of four biological replicates. No significant differences in luciferase transduction were observed between the infected cells.

To better understand the role of the P681H mutation in the maturation and activation of the spike protein, we evaluated the expression and cleavage of the SARS-CoV-2 Wuhan-Hu-1 spike protein (WT), along with the B.1.1.7 and a P681H point mutant of Wuhan-Hu-1 spikes via western blot and densitometry. To do this, we used pseudoparticles consisting of a murine leukemia virus (MLV) core displaying the heterologous viral envelope protein. The pseudoparticles were produced under the normal furin conditions or under the presence of the protease inhibitor decanoyl-RVKR-CMK (dec-RVKR-CMK) to produce cleaved and uncleaved S proteins (respectively). The purified pseudoparticles were later incubated with or without furin, to better assess furin cleavage. As observed in Figure 2B, partial to full cleavage was observed in all the spikes under normal cellular furin conditions (no dec-RVKR-CMK nor exogenous furin), as well as in furin incubated particles. Interestingly, the addition of the dec-RVKR-CMK inhibitor resulted in almost 100% uncleaved spike in WT, but partially cleaved spikes in B.1.1.7 and the P681R mutant. This was also observed when these particles were incubated with exogenous furin, suggesting that the P681H mutation may increase furin cleavage in the complete spike. We then calculated the cleaved S vs. uncleaved S ratio for all these spikes. These data were normalized to the expression of the MLV p30 protein, which is part of the MLV pseudoparticles (Figure 2B). In light of the peptide cleavage data from Figure 2A, it is unclear if this increased cleavage is mediated by furin or another cellular protease, possibly another member of the proprotein convertase (PC) family, of which furin is a member (furin is also defined as *PCSK3*/PC3) (Garten, 2018).

### Functional analysis of virus entry using viral pseudoparticles

To assess the functional importance of the S1/S2 site for SARS-CoV-2 entry, we utilized the MLV pseudoparticles harboring specific SARS-CoV-2 spike proteins. Particles also contain a luciferase reporter that integrates into the host cell genome to drive expression of luciferase, which is quantifiable (Millet et al., 2019). In this study, MLV pseudoparticles containing the SARS-CoV-2 spikes of Wuhan-Hu-1 WT, B.1.1.7, and a P681H point mutant of Wuhan-Hu-1 were also generated alongside positive control particles containing the vesicular stomatitis virus (VSV) G protein, and negative control particles (Δ-Envelope) lacking envelope proteins (not shown), using the HEK293T cell line for particle production.

We examined infection of SARS-CoV-2 pseudoparticles in cell lines representative of both the “early” and “late” cell entry pathways (Figure 2C) as the entry mechanisms of SARS-CoV-2 can be highly cell-type dependent (Whittaker et al., 2021). In this study, we utilized the Vero-TMPRSS2 (“early pathway”) and the Vero-E6 (“late pathway”) cell lines, which are predicted to activate the SARS-CoV-2 S2’ using TMPRSS2 and cathepsin L respectively. While Vero-TMPRSS2 cells gave overall higher luciferase signal indicative of more efficient entry, we observed little difference in infection between pseudoparticles displaying spike protein from either WT, B.1.1.7 or a P681H point mutant. As expected, VSVpp (positive control) infected both cell lines with several orders of magnitude higher luciferase units than the values reported with Δ-Envelope infection (negative control).

### Viral growth in cell culture

According to our results, the P681H mutation did not provide any molecular, nor functional advantage for the B.1.1.7 variant entry into the host cell. However, we wanted to evaluate if the mutations in the B.1.1.7 variant will impact the viral replication of a viral isolate in cell culture (*ex vivo*). To do this, we infected Vero E6 and Vero-TMPRSS2 cells with SARS-CoV-2 strain USA-WA1/2020 or a B.1.1.7 isolate and evaluated the viral growth at 6-, 12-, 24- and 48-hours post-infection (p.i.) through TCID_50_ and immunofluorescence (IFA) assays. We observed that the viral growth of the B.1.1.7 variant was initially slower at 12- and 24-hours p.i., in Vero E6 cells, but it was equivalent to the USA-WA1/2020 growth at 48-hours p.i. (Figure 3A and 3B – Vero E6 cells panels). B.1.1.7 growth in Vero-TMPRSS2 cells was similar to USA-WA1/2020 during the full course of the 48-hours experiment (Figure 3A and 3B – Vero-TMPRSS2 panels). These results agree with our *in vitro* and functional assays, suggesting that the B.1.1.7 variant does not possess a replication advantage to the USA-WA1/2020 strain *ex vivo* in Vero-derived cell lines representative of both the “early” and “late” entry pathways

**Figure 3.**
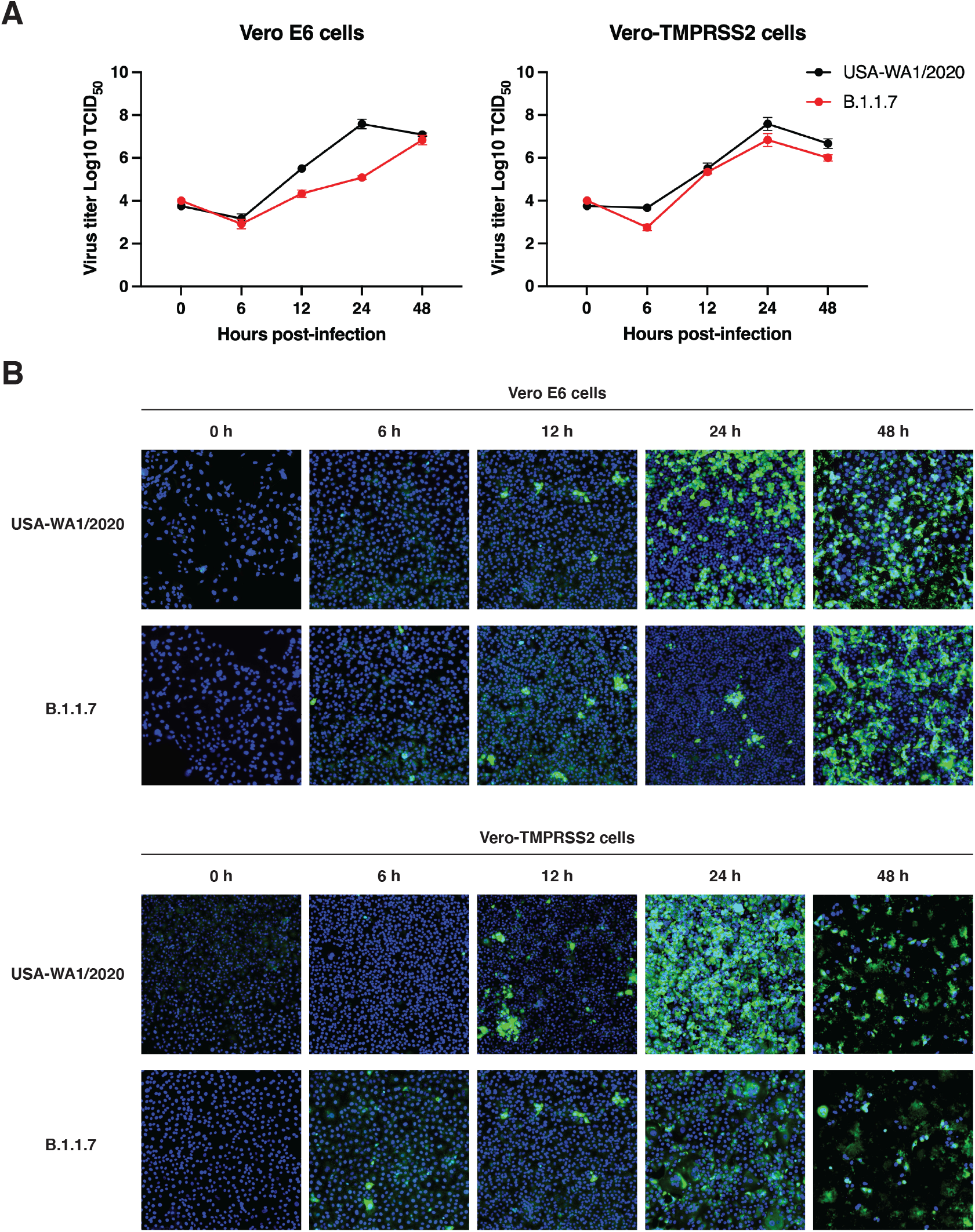
SARS-CoV-2 B.1.1.7 variant infectivity *ex vivo*. **A**. SARS-CoV-2 WA1/2020 and B.1.1.7 growth curves on Vero E6 and Vero-TMPRSS2 cells. Cells were infected to a MOI of 0.1 and supernatant was collected at 0-, 6-, 12-, 24- and 48-hours p.i. Growth curves were then calculated using TCID_50_. **B**. Immunofluorescence assay (IFA) of SARS-CoV-2 WA1/2020 and B.1.1.7 infection on Vero E6 and Vero-TMPRSS2 cells. Cells were infected to a MOI of 0.1 and IFA was performed 0, 6, 12, 24 and 48 hours p.i. Detection of infected cells (green) was conducted using an antibody targeting the SARS-CoV-2 S2 domain. Cell nuclei was stained using DAPI (blue).

### Functional analysis of membrane fusion activity

It has been previously reported that the B.1.1.7 variant possess an advantage in terms of transmission between infected and susceptible individuals (Davies et al., 2021). However, to date, our data does not show functional nor replication features in the B.1.1.7 that support any change in the viral behavior and pathogenesis. Thus, we wanted to explore more directly the fusion capability of the B.1.1.7 spike protein in order to see if the P681H mutation provided any advantage for cell-to-cell transmission. To study this, we performed a cell-to-cell fusion assay where Vero-TMPRSS2 and VeroE6 cells were transfected with the WT, B.1.1.7 and Wuhan-Hu-1 P681H spike gene and we then evaluated syncytia formation as a read-out of membrane fusion (Figure 4A). While Vero-TMPRSS2 cells formed more extensive syncytia than VeroE6 cells, we observed only slight differences in the syncytia formation following spike protein expression for either WT, B.1.1.7, or Wuhan-Hu-1 P681H (Figure 4B). These data show that the P681H mutation had little effect on membrane fusion activity of the SARS-CoV-2 spike protein under the conditions tested.

**Figure 4.**
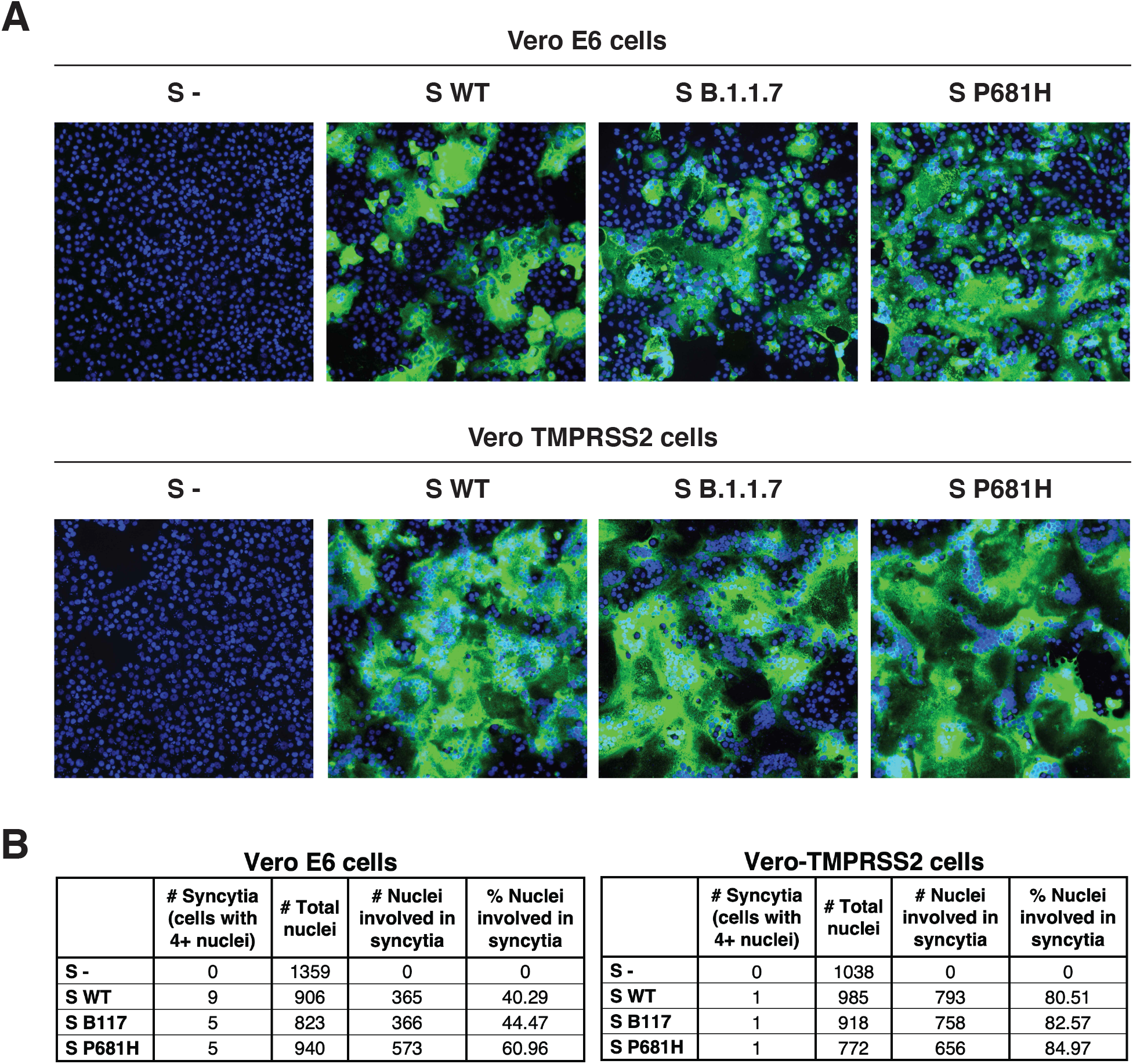
Cell-to-cell fusion in SARS-CoV-2 B.1.1.7 S expressing cells. **A**. Cell-to-cell fusion in Vero E6 and Vero-TMPRSS2 cells expressing SARS-CoV-2 S. Cells were transfected with a plasmid carrying the WT, B.1.1.7, or P681H S gene and evaluated through IFA after 28 hours. Syncytia formation was observed using an antibody targeting the SARS-CoV-2 S2 domain and a DAPI for cell nuclei (blue). **B**. Syncytia counting in SARS-CoV-2 S expressing cells. Total of nuclei per cell were counted to determine the percentage of nuclei involved in syncytia.

### SARS-CoV-2 B.1.1.7 variant infection in human respiratory tract-derived cells

So far, all our results have shown no differences between B.1.1.7 and WT behavior from molecular and functional points of view. However, the functional studies were performed using susceptible cells that do not necessarily represent the typical cell type that the virus will initially infect *in vivo*, which are principally respiratory tract cells. To address this, we performed functional viral entry studies and live virus infection assays in respiratory-type Calu-3 cells. We used the MLV pseudoparticles harboring the WT, B.1.1.7 and mutated P681H spike proteins and infected Calu-3 cells. We observed slight but significant differences in the luciferase transduction between WT and B.1.1.7 spike-carrying pseudoparticles (Figure 5A), but no differences with the P681H mutated spike-carrying particles., compared to WT. Interestingly, we did not observe any differences in the viral growth between USA-WA1/2020 and a B.1.1.7 virus isolate in Calu-3 cells (Figure 5B), which suggests that, as with Vero E6 and Vero-TMPRSS2 cells, there no functional advantage for B.1.1.7 in respiratory epithelial cells; i.e. Calu-3 cells.

**Figure 5.**
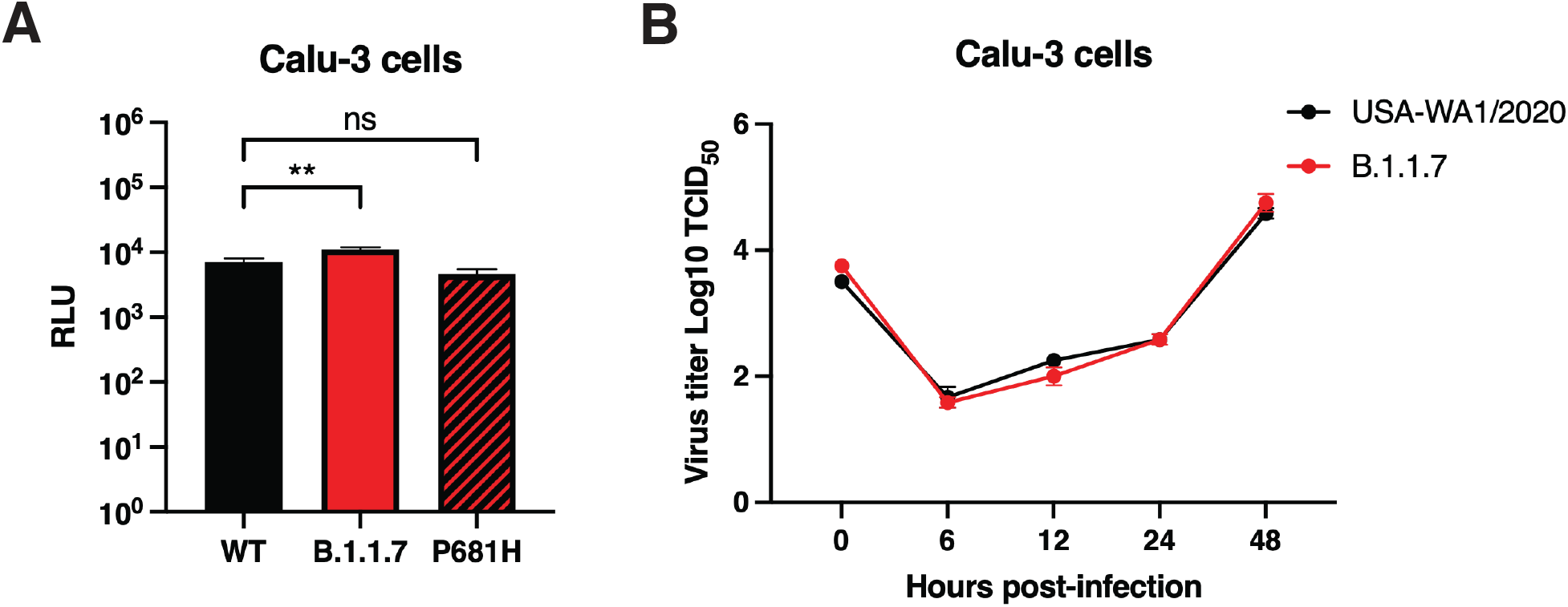
Infection in respiratory Calu-3 cells. **A**. Pseudoparticle infectivity assays in Calu-3 cells. Cells were infected with MLV pseudoparticles harboring the VSV-G, SARS-CoV-2 S WT, SARS-CoV-2 S B.1.1.7 variant, SARS-CoV-2 S WT with P681H mutation. Data represents the average luciferase activity of cells of three biological replicates. A significant increase in the luciferase transduction were observed with the pseudoparticles harboring the B.1.1.7 S (P=0.0076). **B**. SARS-CoV-2 WA1/2020 and B.1.1.7 growth curves on Calu-3 cells. Cells were infected to a MOI of 0.1 and supernatant was collected at 0-, 6-, 12-, 24- and 48-hours p.i. Growth curves were then calculated using TCID_50_.

## Discussion

The factors influencing increased transmissibility and pathogenicity of SARS-CoV-2 variants remain poorly understood. Here, we performed *in vitro* assays using fluorogenic peptides mimicking the S1/S2 sequence from both Wuhan-Hu-1 and the B.1.1.7 VOC and observed no definitive difference in furin cleavage for B.1.1.7. We performed functional assays using pseudo-typed particles harboring SARS-CoV-2 spike proteins and observed no significant transduction differences between Wuhan-Hu-1 and B.1.1.7 spike-carrying pseudo-typed particles in VeroE6 or Vero-TMPRSS2 cells, despite the spikes containing P681H being more efficiently cleaved. Likewise, we show no differences in cell-cell fusion assays using the spike P681H-expressing cells, as well as no notable effects on viral replication. Our findings suggest that the introduction of P681H in the B.1.1.7 variant may increase spike cleavage by furin-like proteases, but this does not significantly impact viral entry, infection or cell-cell spread. We consider that other factors are at play to account for the increased in transmission and disease severity attributed to the SARS-CoV-2 B.1.1.7 VOC.

Overall, we show that the P681H mutation at the S1/S2 site of the SARS-CoV-2 spike protein may increase its cleavability by furin-like proteases, but that this does not translate into increased virus entry or membrane fusion. These findings are broadly in line with those of Brown *et al*. (Brown et al., 2021) using infectious SARS-CoV-2, who showed no differences in the B.1.1.7 variant in terms of viral replication in primary human airway cells but did show a disadvantage in Vero cells linked to increased cleavage of the spike protein. In contrast, Dicken *et al*. (Dicken et al., 2021) indicated enhanced entry of B.1.1.7, but in this case only under conditions with low expression of the ACE2 receptor.

B.1.1.7 is certainly not the only SARS-CoV-2 variant with a P681H change in the spike protein; it is also present in B.1.243 (clade 20A), B.1.222 (clade 20B) and a lineage B.1 variant termed clade 20C, with these three variants recently reported in New York State, USA (Lasek-Nesselquist et al., 2021). Interestingly, B.1.243 comprised the majority of P681H-containing viruses and was the predominant variant in New York in November 2020, but had declined significantly by February 2021 (to be replaced by B.1.1.7 and B.1.222 among other variants) some of which do not contain P681H (e.g., B.1.429). Other examples of variants containing P681H include A.VOI.V2 detected through travel surveillance in Angola, Africa (de Oliveira et al., 2021), isolates from Hawaii (Maison et al., 2021) and viruses originally classified under lineage B.1.1.28 in locations such as the Philippines (Tablizo et al., 2021). However, many other VOIs e.g., CAL.20C (Zhang et al., 2021) do not contain P681H. Other variants containing mutations have been identified that have been proposed to impact S1/S2 cleavage, e.g., A688V (Tegally et al., 2021b), but we consider such mutations to be too distal to the cleavage site to have a direct impact on furin activity.

The SARS-CoV-2 S1/S2 site contains three basic residues in atypical spacing for a furin cleavage site (Tang et al., 2021), and as such is not “polybasic”. The P681H mutation increases the number of basic residues to four (especially at lowered pH) and is predicted to result in a slightly increased cleavability based on ProP and Pitou scoring and peptide cleavage assays at pH 7.4 (Figure 1A). However, it is important to note that the P681H change does not result in the formation of a consensus furin cleavage site (i.e., ^P4^**R**-X-**K/R**-**R**^P1^), even at lowered pH. It is also of note that increased spike protein cleavage does not always translate into increased function and may in fact be detrimental. A good example of this is the insertion of furin cleavage sites into SARS-CoV-1 spike at S1/S2 and/or S2’, which resulted in a hyper-fusogenic phenotype (Belouzard et al., 2009; Follis et al., 2006), but with pseudoparticles being unrecoverable especially when added at S2’ presumably due to the presence of an unstable spike protein. One interpretation of the appearance of the P681H mutation in circulating viruses is that SARS-CoV-2 is evolving to become a transmissible but relatively benign community-acquired respiratory (CAR) or “common cold” coronavirus (Jin et al., 2020), such as the betacoronaviruses HCoV-HKU1 and HCoV-OC43 both of which have very strong “polybasic” furin cleavage sites at the S1/S2 position (Figure 1A). It remains to be determined how any increased cleavage of SARS-CoV-2 S may affect factors such as increased duration of viral shedding, which is one possible explanation of increased transmissibility for B.1.1.7. (Kissler et al., 2021).

As explained for other variants (including D614G in humans and Y453F in mink (Lauring and Hodcroft, 2021), we consider that the rise of P681H-containing viruses could be due to chance, with founder effects being responsible for rapid progression, or P681H may confer an advantage with regards transmissibility or cell-cell spread *in vivo*. Our data here reinforce the concept that while analysis of individual point mutations is critical part of understanding virus biology, a full assessment of the epidemiological context of virus infection requires more extensive study (Goodman and Whittaker, 2021). It will continue to be important to track VOCs and other variants that may pose a threat for exponential spreading. While the VOCs B.1.351 and B.1.1.28.1 (P.1) have been of high concern due to immune escape (Coutinho et al., 2021; Darby and Hiscox, 2021; Garcia-Beltran et al., 2021; Sabino et al., 2021; Tegally et al., 2020), B.1.1.7 was not considered a concern in this regard (Planas et al., 2021); however, the possible acquisition of “immune-escape” mutations such as E484K or N439K (Chan et al., 2020; Di Caro et al., 2021) remains possible. While B.1.1.7 remains in circulation, B.1.617.2 is now outcompeting this and other variants. The three notable B.1.617 variants B.1.617.1, B.1.617.2 (Delta) and B.1.617.3, along with prior variants such as A.23.1 all contain a distinct change in the P5 position of the S1/S2 cleavage site (P681R), with early indications suggesting a growth advantage over P681H (Peacock et al., 2021; Saito et al., 2021), possibly with the replacement of the histidine in B.1.1.7 with a more conventional basic amino acid providing an important increase in viral fitness in the pathway of pandemic progression. We have recently addressed the role of the P681R mutation on the activation by furin, and found that the introduction of an arginine at this position, could significantly increase furin activation, potentially increasing viral entry and cell-to-cell spread (Lubinski et al., 2021). Other recent studies have also demonstrated that the P681R mutation enhances the viral replication, and the introduction of a reverse mutation to the WT genotype P681 significantly reduces the fitness of the Delta variant, to levels even lower than the Alpha variant, which harbors the P681H mutation (Liu et al., 2021). These findings highlight the importance of the P681R mutation and provide insights in the increased fitness of B.1.617.2/Delta.

B.1.1.7 remains as a VOC as defined by the World Health Organization, and has recently been moved to a “variant being monitored” (VBM) as defined by the US Centers for Disease Control (CDC). Despite the current global dominance of B.1.617.2/Delta (containing P618R), other P681H-containing variants such as B.1.620 (Dudas et al., 2021) contain an even higher level of spike mutations than B.1.1.7 (11 in total, including P681H) and are expanding in certain regions of Europe and Africa, which means that the evolution of SARS-CoV-2 remains on ongoing process that needs to be carefully assessed.

## Limitations of the study

Bioinformatic and *in silico* analysis are predictive and they may not be fully accurate. These tools provided the bases for the subsequent experiments, but they do not provide definite data. We used fluorogenic peptides mimicking the SARS-CoV-2 S1/S2 regions, to evaluate the cleavage by furin and trypsin proteases. These short-length peptides do not resemble the original structural folding in the native protein, and this could result in an altered cleavage by the protease. Experiments using purified full-length protein may be needed to further evaluate the protease cleavage. The pseudoparticle system we used for the viral entry functional experiments, has been broadly described and used in previous studies by our group and other researchers globally. This system provides a safe tool to study high pathogenic viruses under biosecurity level 2 (BSL-2) conditions, as the pseudoparticles can mimic the viral entry but their replication is impaired, which eliminates the biosafety risks. While lentivirus based pseudoparticles (like the MLV used by us) are commonly used to study coronaviruses, these systems have the disadvantage that their assembly and viral envelope protein incorporation differs from the normal pathways used by coronaviruses. This may affect the amount of viral envelope protein that is incorporated to the pseudoparticles and hindering the evaluation of the viral infection. We performed live virus infection assays to corroborate our pseudoparticle data, and observed a similar behavior, which suggests that the pseudoparticle system is accurate and valid, despite differential membrane envelope protein incorporation between MLV and coronaviruses.

## Acknowledgements

This work was funded by the National Institute of Health research grant R01AI35270. TT is supported by the National Science Foundation Graduate Research Fellowship Program under Grant No. DGE-1650441 and the Samuel C. Fleming Family Graduate Fellowship. We would especially like to thank Hector Aguilar-Carreno and Ruth Collins for important insight, and all members of the Daniel, Diel and Whittaker groups for helpful discussions.

## Author contributions

Conceptualization: B.L., T.T., S.D., J.A.J. and G.R.W.; Methodology: B.L., M.H.V.F., T.T., S.D., D.D., J.A.J. and G.R.W; Investigation: B.L., M.H.V.F., T.T., J.A.J.; Writing - Original Draft: B.L., T.T., J.A.J. and G.R.W; Writing - Review & Editing, B.L., M.H.V.F., T.T., S.D., D.D., J.A.J. and G.R.W; Visualization: B.L., M.H.V.F., T.T. and J.A.J.; Supervision: S.D., D.D., J.A.J. and G.R.W.; Funding acquisition: S.D. and G.R.W.

## Methods

### Furin prediction calculations

Prop: CoV sequences were analyzed using the ProP 1.0 Server hosted at: cbs.dtu.dk/services/ProP/. PiTou: CoV sequences were analyzed using the PiTou V3 software hosted at: http://www.nuolan.net/reference.html.

### *In silico* homology modelling

SARS-CoV-2 S protein models for Wuhan Hu-1 (GenBank accession # MN908947.3) and B.1.1.7 hCoV-19/England/MILK-9E05B3/2020 (GISAID accession # EPI_ISL_601443 – Original sample and sequence submission information included in table 1), were built using UCSF Chimera (v.1.14, University of California) through the homology modeling tool of the Modeller extension (v.10.1, University of California) as described in (Jaimes et al., 2020a). Models were built based on the SARS-CoV S structure (PDB No. 5×58).

**Table 1:** SARS-CoV-2 B.1.1.7 variant original sample and sequence submission information.

| Virus Name | Accession Number | Originating Laboratory | Submitting Laboratory | Authors | Submitter |
| --- | --- | --- | --- | --- | --- |
| hCoV-19/England/MILK-9E05B3/2020 | EPI_ISL_601443 | Lighthouse Lab in Milton Keynes | Wellcome Sanger Institute for the COVID-19 Genomics UK (COG-UK) consortium | The Lighthouse Lab in Milton Keynes and Alex Alderton, Roberto Amato, Sonia Goncalves, Ewan Harrison, David K. Jackson, Ian Johnston, Dominic Kwiatkowski, Cordelia Langford, John Sillitoe on behalf of the Wellcome Sanger Institute COVID-19 Surveillance Team (http://www.sanger.ac.uk/covid-team) | Robert Davies |

### Fluorogenic peptide assays

Fluorogenic peptide assays were performed as described previously with minor modifications (Jaimes et al., 2019). Each reaction was performed in a 100 μL volume consisting of buffer, protease, and SARS-CoV-2 S1/S2 WT (TNSPRRARSVA) or SARS-CoV-2 S1/S2 B.1.1.7 (TNSHRRARSVA) fluorogenic peptide in an opaque 96-well plate. For trypsin catalyzed reactions, 0.8 nM/well TPCK trypsin was diluted in PBS buffer. For furin catalyzed reactions, 1 U/well recombinant furin was diluted in buffer consisting of 20 mM HEPES, 0.2 mM CaCl2, and 0.2 mM β-mercaptoethanol, at pH 6.5, 7.0 or 7.5. Fluorescence emission was measured once per minute for 60 continued minutes using a SpectraMax fluorometer (Molecular Devices, Inc.) at 30 °C with an excitation wavelength of 330 nm and an emission wavelength of 390 nm. Vmax was calculated by fitting the linear rise in fluorescence to the equation of a line.

### Synthesis and cloning of the B.1.1.7 spike protein

The B.1.1.7 spike gene from isolate hCoV-19/England/MILK-9E05B3/2020 (EPI_ISL_601443) was codon-optimized, synthesized and cloned into a pcDNA 3.1+ vector for expression (GenScript Biotech Co.).

### Site-directed mutagenesis

Mutagenesis primers (cagacctggctctcctgtgggagtttgtctgggt/ acccagacaaactcccacaggagagccaggtctg) were designed based on the DNA sequence for SARS-CoV-2 Wuhan-Hu-1 using the QuickChange Primer Design tool (Agilent Technologies, Inc.). Mutagenesis was carried out on a pCDNA-SARs2 Wuhan-Hu 1 S plasmid to create the P681H mutation, using the QuickChange Lightning Mutagenesis kit (Agilent Technologies, Inc.). The original plasmid was generously provided by David Veesler, University of Washington USA. XL-10 gold competent cells were transformed with the mutated plasmid, plated on LB Agar + Ampicillin plates, and left at 37°C overnight. A distinct colony was chosen the next day to grow up a 4 ml small culture at 37°C overnight. pCDNA-SARC-CoV-2 Wuhan-Hu-1 P681H S plasmid was then extracted using the QIAprep Spin Miniprep Kit (Qiagen N.V.) and Sanger sequencing was used to confirm incorporation of the mutation.

### Pseudoparticle generation

HEK-293T cells were seeded at 3×10^5^ cells/ml in a 6-well plate the day before transfection. Transfection was performed using polyethylenimine (PEI) and 1X Gibco^®^ Opti-Mem (Life Technologies Co.). Cells were transfected with 800ng of pCMV-MLV *gag-pol*, 600ng of pTG-Luc, and 600 ng of a plasmid containing the viral envelope protein of choice. Viral envelope plasmids included pCAGGS-VSV G as a positive control, pCDNA-SARS-CoV-2 Wuhan-Hu-1 S, pCDNA-SARS-CoV-2 (Wuhan-Hu-1) P681H S, and pCDNA-SARS-CoV-2 B.1.1.7 S. pCAGGS empty vector was used for a Δ-Envelope negative control. 48 hours post transfection, the supernatant containing the pseudoparticles was removed, centrifuged to remove cell debris, filtered, and stored at -80°C. For + dec-RVKR-CMK particles, 7.5 μL of dec-RVKR-CMK was added to cells immediately after transfection.

### Pseudoparticle Infection Assay

Vero E6 and Vero-TMPRSS2 cells were seeded at 3×10^5^ cells/ml in a 24 well plate the day before infection. Cells were washed three times with 1X DPBS and then infected with 200 µl of either VSV G, SARS-CoV-2 S, SARS-Cov-2 P681H S, SARS-CoV-2 B.1.1.7 S, or Δ-Envelope pseudoparticles. Infected cells incubated on a rocker for 1.5 hours at 37°C, then 300 µl of complete media were added and cells were left at 37°C. At 72 hours post-infection, cells were lysed and the level of infection was assessed using the Luciferase Assay System (Cat: E1501, Promega Co.). The manufacturer’s protocol was modified by putting the cells through 3 freeze/thaw cycles after the addition of 100 µl of the lysis reagent. 10 µl of the cell lysate were added to 20 µl of luciferin, and then luciferase activity was measured using the Glomax 20/20 luminometer (Promega Co.). Infection assays were done in triplicate and were replicated 4 times. All four replicates were carried out using pseudoparticles generated from the same transfection.

### Western blot analysis of pseudoparticles

3 mL of pseudoparticles were pelleted using a TLA-55 rotor with an Optima-MAX-E ultracentrifuge (Beckman Coulter, Inc.) for 2 hours at 42,000 rpm at 4°C. Untreated particles were resuspended in 30 µL DPBS buffer. For the + furin treated MLVpps, particles were resuspended in 30 μL of furin buffer consistent in 20 mM HEPES, 0.2 mM CaCl2, and 0.2 mM β-mercaptoethanol (at pH 7.0). Particles were later incubated with 6 U of recombinant furin for 3 h at 37 °C. Sodium dodecyl sulfate (SDS) loading buffer and DTT were added to samples and heated at 65°C for 20 minutes. Samples were separated on NuPAGE Bis-Tris gel (Cat: NP0321BOX, Invitrogen Co.) and transferred on polyvinylidene difluoride (PVDF) membranes (MilliporeSigma). SARS-CoV-2 S was detected using a rabbit polyclonal antibody against the S2 domain (Cat: 40590-T62, Sino Biological, Inc) and an AlexaFluor™ 488 goat anti-rabbit antibody (Invitrogen Co, USA). Bands were detected using the ChemiDoc (Bio-Rad Laboratories, Inc.) and band intensity was calculated using the analysis tools on Image Lab 6.1 software (Bio-Rad Laboratories, Inc.) to determine the uncleaved-to-cleaved S ratios.

### Growth curves

SARS-CoV-2 WA1/2020 and B.1.1.7 growth curves were performed in Vero E6, Vero-TMPRSS2 and Calu-3 cells. Cells were cultured in 24-well plates in advance, inoculated with a multiplicity of infection (MOI) of 0.1 and harvested at 6-, 12-, 24- and 48-hours post-infection (p.i.) in triplicate. Virus titers were determined on each time by limiting dilutions in Vero E6 cells. At 48-hours p.i. cells were fixed (3.7% formaldehyde), permeabilized (0.2% Triton-X), and stained with an anti-NP SARS-CoV-2 rabbit polyclonal antibody (produced and characterized in Diel laboratory) and then incubated with a goat anti-rabbit IgG secondary antibody (Alexa Fluor 594; Immunoreagents). Viral titers were determined by the Spearman and Karber’s method and expressed as tissue culture infectious dose 50 (TCID_50_) per milliliter.

### Immunofluorescence assay

Vero E6 and Vero-TMPRSS2 cells cultured in 4-well glass slides were infected with SARS-CoV-2 WA1/2020 and B.1.1.7 (MOI = 0.1). At 0-, 6-, 12-, 24- and 48-hours p.i. cells were fixed using 3.7% formaldehyde in PBS (pH 7.2) for 30 min. The cells were washed three times with PBS and quenched with 50 mM NH4Cl. Permeabilization was performed with 0.1% Triton X-100 in PBS for 5 minutes on ice. Blocking was performed using 5% heat inactivated goat serum in PBS for 20 minutes and the antibodies for labeling were diluted in the same solution. The spike expression was detected using the SARS-CoV-2 spike antibody (Cat: 40591-T62, Sino Biological Inc.) at 1/500 dilution for 1 hour. Secondary antibody labeling was performed using AlexaFluor™ 488 goat anti-rabbit IgG antibody (Cat: A32731, Invitrogen Co.) at a 1/500 dilution for 45 minutes. Three washes with PBS were performed between each step of the assay. Finally, slides were mounted using DAPI Fluoromount-G^®^ (Cat: 0100-20, SouthernBiotech Inc.) and analyzed with fluorescence.

### Cell-cell fusion assay

Vero E6 and Vero-TMPRSS2 cells were transfected with a plasmid harboring the spike gene of the SARS-CoV-2 Wuhan-Hu 1, SARS-CoV-2 B.1.1.7, the SARS-CoV-2 P681H S, or a delta-spike pCDNA3.1+ plasmid, and evaluated through an immunofluorescence assay (IFA). Transfection was performed on 8-well glass slides at 90% confluent cells using Lipofectamine^®^ 3000 (Cat: L3000075, Invitrogen Co.), following the manufacturer’s instructions and a total of 250 ng of DNA per well was transfected. The cells were then incubated at 37°C with 5% of CO_2_ for 28 hours. Syncytia was visualized through an immunofluorescence assay using the method described in the previous section. Representative images of each treatment group were taken, and these images were used to calculate the percent of nuclei involved in the formation of syncytia. Images were taken at 20X on the Echo Revolve fluorescent microscope (Model: RVL-100-M). The nuclei were counted manually using the Cell Counter plugin in ImageJ (https://imagej.nih.gov/ij/). Cells that expressed the spike protein and contained 4 or more nuclei were considered to be one syncytia.

